# A Genomic Reference Panel for *Drosophila serrata*

**DOI:** 10.1101/266031

**Authors:** Adam R. Reddiex, Scott L. Allen, Stephen F. Chenoweth

## Abstract

Here we describe a collection of re-sequenced inbred lines of *Drosophila serrata*, sampled from a natural population situated deep within the species endemic distribution in Brisbane, Australia. *D. serrata* is a member of the speciose *montium* group whose members inhabit much of south east Asia and has been well studied for aspects of climatic adaptation, sexual selection, sexual dimorphism, and mate recognition. We sequenced 110 lines that were inbred via 17-20 generations of full-sib mating at an average coverage of 23.5x with paired-end Illumina reads. 15,228,692 biallelic SNPs passed quality control after being called using the Joint Genotyper for Inbred Lines (JGIL). Inbreeding was highly effective and the average levels of residual heterozygosity (0.86%) were well below theoretical expectations. As expected, linkage disequilibrium decayed rapidly, with r^2^ dropping below 0.1 within 100 base pairs. With the exception of four closely related pairs of lines which may have been due to technical errors, there was no statistical support for population substructure. Consistent with other endemic populations of other *Drosophila* species, preliminary population genetic analyses revealed high nucleotide diversity and, on average, negative Tajima’s D values. A preliminary GWAS was performed on a cuticular hydrocarbon trait, 2-MeC_28_ revealing 4 SNPs passing Bonferroni significance residing in or near genes. One gene *Cht9* may be involved in the transport of CHCs from the site of production (oenocytes) to the cuticle. Our panel will facilitate broader population genomic and quantitative genetic studies of this species and serve as an important complement to existing *D. melanogaster* panels that can be used to test for the conservation of genetic architectures across the *Drosophila* genus.

## Introduction

The availability of whole genome sequence data for *Drosophila* species has greatly facilitated advances in the fields of genetics and evolutionary biology. For example, the sequencing of 12 *Drosophila* genomes (Clark et al. 2007) was instrumental to new discoveries in comparative genomics (Stark et al. 2007; Sturgill et al. 2007; Zhang et al. 2007). The advent of affordable genome sequencing has also allowed population geneticists to characterise genomic variation within and among natural populations, improving our understanding of the complex evolutionary histories of cosmopolitan species such as *D. melanogaster* and *D. simulans* (Begun et al. 2007; Lack et al. 2015; Langley et al. 2012; Pool et al. 2012). Most recently, multiple panels of re-sequenced inbred *D. melanogaster* lines have become available, facilitating the molecular dissection of complex trait variation (Grenier et al. 2015; Huang et al. 2014; King et al. 2012; Mackay et al. 2012). With these populations of reproducible genotypes, researchers have used genome-wide association analysis to identify genetic variants underlying variation in a broad range of traits including physiological traits (Burke et al. 2014; Dembeck et al. 2015; Gerken et al. 2015; Unckless et al. 2015; Weber et al. 2012), behaviours (Shorter et al. 2015), recombination rates (Hunter et al. 2016), disease susceptibility (Magwire et al. 2012), and traits related to human health (Harbison et al. 2013; He et al. 2014; King et al. 2014; Kislukhin et al. 2013; Marriage et al. 2014).

Just as comparative genomic and population genetic studies of adaptation (e.g. Machado et al. 2016; Zhao et al. 2015) have been enhanced through the availability of multi-species genome resources, quantitative genetics may also benefit from the availability of multispecies genome panels. The development of panels of re-sequenced lines for *Drosophila* species beyond *D. melanogaster* may support broader lines of inquiry such as the conservation of genetic architectures among related taxa (Yassin et al. 2016). To this end, we have developed a new genomic resource for *D. serrata*, a member of the *montium* group of species. The *montium* group has long been regarded as a subgroup within the melanogaster species group (Lemeunier et al. 1986), but has more recently been considered as a species group of its own (Da Lage et al. 2007; Yassin 2013). Although *montium* contains 98 species (Brake and Bachli 2008) and represents a significant fraction of all known *Drosophila* species, there have been very few genomic investigations of its members. Recently, genomic tools have been developed for *D. serrata* including an expressed sequence tag (EST) library (Frentiu et al. 2009), a physical linkage map (Stocker et al. 2012), and transcriptome-wide gene expression datasets (Allen et al. 2013; Allen et al. 2017a; McGuigan et al. 2014). Additionally, an assembled and annotated genome sequence (Allen et al. 2017b) make *D. serrata* only the second species in the *montium* group with a sequenced genome after *D. kikkawai* (NCBI *Drosophila kikkawai* Annotation Release 101). Coupled to this, *D. serrata* is one member of the *montium* group that has been extensively studied in the field of evolutionary genetics.

Populations of *D. serrata* have been recorded from as far north as Rabaul, Papua New Guinea (4.4°N) (Ayala 1965) to as far south as Woolongong, Australia (34.3°S) (Jenkins and Hoffmann 1999). This broad latitudinal range has made *D. serrata* an ideal model for population studies addressing the evolution of species borders (Blows and Hoffmann 1993; Hallas et al. 2002; Magiafoglou et al. 2002; van Heerwaarden et al. 2009) and adaptation along latitudinal clines (Allen et al. 2017a; Frentiu and Chenoweth 2010; Kellermann et al. 2009). *D. serrata* has also emerged as a powerful model for the application of quantitative genetic designs to investigate sexual selection (Gosden and Chenoweth 2014; Hine et al. 2002; McGuigan et al. 2011).

Here, we the report development of a panel of 110 re-sequenced inbred *D. serrata* lines that we have called the *Drosophila serrata* Genome Reference Panel (DsGRP). Similar to the DGRP (Mackay et al. 2012), flies were sampled from a single large natural population with the exception that *D. serrata* was sampled from its endemic distribution. In this initial description, we estimate the degree of heterozygosity remaining in the lines after inbreeding, show the degree to which lines are genetically related to one another, estimate genome-wide levels of nucleotide diversity, and describe patterns of linkage disequilibrium. We also demonstrate how this panel of flies can be used to genetically dissect trait variation by performing a genome-wide association analysis on variation in a cuticular hydrocarbon (CHC) trait.

## Methods

### Collection and inbreeding

*Drosophila serrata* were collected from a wild population located at Bowman Park, Brisbane Australia (Latitude: -27.45922, Longitude: 152.97768) during October 2011. We established each line from a single, gravid female before applying 20 generations of inbreeding. Inbreeding was carried out each generation by pairing virgin brothers and sisters. 100 inbred lines, out of the initial 239 iso-female lines established, survived the full 20 generations inbreeding and a further 10 lines were established after 17 generations of inbreeding.

### Sequencing

We sequenced the genomes of 110 inbred lines using 100 base-pair paired-end reads with a 500 base-pair insert on an Illumina Hiseq 2000 sequencing machine. Sequencing and library preparation were carried out by the Beijing Genomics Institute. DNA from each line was isolated from a pool of at least 30 virgin female flies using a standard phenol-chloroform extraction method.

### Quality control and SNP calling

We received reads from the Beijing Genomics Institute for which approximately 95% of the bases from each line had a base quality score greater than or equal to 20 (Illumina GA Pipeline v1.5). Read quality was also assessed using FastQC v0.11.2 before being mapped to the *Drosophila serrata* reference genome (Allen et al. 2017b) using BWA-mem v0.7.10 (Li 2013) and were realigned around indels using the GATK IndelRealigner v3.2-2 (McKenna et al. 2010). Genotypes for every line were inferred simultaneously using the Joint Genotyper for Inbred Lines (JGIL) v1.6 (Stone 2012). This probabilistic model was especially designed for genotyping large panels of inbred lines or strains and is considered to have high accuracy (Mackay et al. 2012; Stone 2012). Genotype calls with a probability lower than 99% were treated as missing genotypes.

### Residual heterozygosity

The residual heterozygosity per line was estimated as the genome-wide proportion of sites that remained heterozygous after 17-20 generations of inbreeding, more specifically, we summed all of the genotype call that were heterozygous and expressed this statistic as a percentage of all genotyped sites. In addition, for each site in the genome that differed among the inbred lines, we calculated the percentage of lines that were heterozygous for that site. Site filtering based on minor allele frequency and coverage was not performed for this analysis.

### Pairwise relatedness between lines

Pairwise relatedness between lines (*j* and *k*) was estimated using the ‑‑make-grm-inbred command of GCTA v1.24.2 (Yang et al. 2011), which applies the expression:

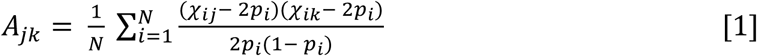

where, *χ_ij_* is the number of copies of the reference allele for the *i*^th^ SNP for individual *j* and *p* is the population allele frequency. *N* is the total number SNPs. Only biallelic SNPs with a read depth between 5 and 60 and a minor allele frequency above 5% were used to estimate relatedness.

### Estimating population substructure

To test whether the sample of lines exhibited any underlying substructure, we used the approach of Bryc et al. (2013) which is founded on random matrix theory. Importantly, while the genomic relatedness matrix calculated in GCTA, **A** = **WW’**/*N*, is normalised with element 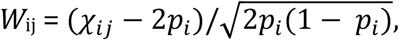 is the number copies of the minor allele carried by individual *j* (Yang et al. 2011) and *p_i_* is the population allele frequency, the genomic relatedness matrix used by the approach of Bryc et al. (2013) is not. For this approach, the genomic relatedness matrix, **X** = **CC’**, where **C** is an M x N rectangular matrix with M corresponding to the number of individuals used to estimate **X** and N is the number of SNPs, here the element *C*_ij_ = *χ_ij_*, the number of copies of the minor allele carried by individual *j*. **X** was scaled by equation 2.8 in Bryc et al. (2013) using values of M= 110 and N=3,709,328. Only SNPs without missing data were used for this analysis. The number of sub-populations was determined by the number of eigenvalues larger than that expected for a random relatedness matrix given the significance threshold *t* = (1 + *F*)/2 (Bryc et al. 2013). Here, *t* corresponds to a value of 0.993 with our expected inbreeding coefficient (*F*) of 0.986 (Falconer and Mackay 1996) after 20 generations of full-sib mating. Upon seeing that a small number of lines were unusually highly related (see results), we repeated this analysis after removing four lines (line IDs: 29, 134, 159, 206) to verify that the significant results were driven only by these “outliers”.

### Linkage disequilibrium

We estimated linkage disequilibrium as the square of the inter-variant allele count correlation (r^2^) using PLINK v1.9 (Chang et al. 2015). We estimated r^2^ in non-overlapping 500 base-pair windows across the entire genome. The analysis was performed on biallelic SNPs that had an average read depth of between 5 and 60 and a minimum minor allele frequency of 5%. The four highly related lines (r >= 0.1) were removed prior to the analysis.

### Nucleotide diversity and neutrality

We estimated nucleotide diversity (π) and Tajima’s D (Tajima 1989) statistic in 50 kilobase non-overlapping sliding windows and took the mean for each of the major chromosome arms (2L, 2R, 3L, 3R, and X) using vcftools v0.1.15 (Danecek et al. 2011). The SNP data used for this analysis had an average read depth across all lines of between 5 and 60 but no threshold on minor allele frequency was applied. Again, the four highly related lines were removed prior to this analysis.

### Genome-wide association of female CHC expression

As proof-of-concept we performed a GWAS on a single cuticular hydrocarbon, 2-methyloctacosane, (2-Me-C_28_). CHCs are waxy substances that are secreted on the cuticle with 2-Me-C_28_ being one of a suite of CHCs that have been extensively studied in this species due to their role in species recognition, mate choice, and desiccation resistance. For each of 94 lines, we extracted CHCs from four virgin females, across two replicate vials using individual whole-body washes in 100µl of the solvent hexane. We used a standard gas chromatography method to quantify the amount of 2-Me-C_28_ (Blows and Allan 1998). To maintain the trait scale used in previous studies, we transformed the amount of 2-Me-C_28_ into a log-contrast value following Aitchison (1986), using an additional trait, the CHC 9-hexacosane, 9-C_26:1_, as the divisor. This transformation turns the expression of CHCs into a proportional measure and provides an internal control for other sources of variation including body size and condition.

Our GWAS contrasts with other analyses performed on the DGRP in that we model trait variation at the individual, rather than line mean level. We applied a single marker mixed effects association analysis, where the following model was fit for every biallelic SNP that had mean coverage between 5 and 60 and sample MAF of 5%:

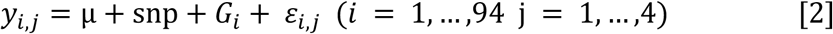

Here, CHC expression (y) of replicate individual *j* on genotype *i* is modelled as a function of the mean term (μ), the additive fixed effect of the candidate SNP, the polygenic random effect that is captured by the genomic relatedness matrix (***G***) where ***G*** has a 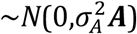 distribution, and the residual error (*ε*). This model was specifically designed for populations where identical genotypes can be measured independently in multiple organisms such as inbred lines (Kruijer et al. 2015) and was fit using AsReml-R v3 (VSN International) for a total of 3,318,503 SNPs. There are a couple of differences between the approach outlined above and other mixed modelling approaches to GWAS implemented in programs such as GEMMA (Zhou and Stephens 2012), FaST-LMM (Lippert et al. 2011), and GCTA (Yang et al. 2011). First, the use of individual-level opposed to line mean level observations, allows for estimation of the genomic heritability (Kruijer et al. 2015). Although mapping power is unlikely to be significantly boosted through the use of individual level data *per se* when working with inbred lines (Kruijer et al. 2015), a second aspect to our approach does potentially increase power. Our, albeit slower, approach re-estimates the polygenic variance component when each SNP tested and results in an exact calculation of Wald’s test statistic (Zhou and Stephens 2012). This contrasts with the approach used by many mixed model GWAS programs where, to save computation time, this variance component is estimated once in a null model with no fixed effect of SNP and then held constant for each SNP tested. Such an approach produces an approximate value of the test statistic which can result in power loss under some circumstances (Zhou and Stephens 2012). As these circumstances are difficult to predict beforehand, we chose to re-estimate the polygenic random effect despite the computational cost.

To increase computational speed, we nested this linear model within an R loop that allows the access of multiple cores using the “foreach” and “doMC” packages (Revolution Analytics and Steve Weston 2015). Significant SNPs were identified as those with p-values that passed Bonferroni multiple test correction –log10(p) > 7.8, we also report SNPs with –log10(p) > 5 for comparison to other *Drosophila* GWAS where this arbitrary threshold value is used (Mackay et al. 2012). We took statistically significant SNPs and annotated them to the current version of the *D. serrata* genome (Allen et al. 2017b). If a significant SNP was located within a gene, we blasted the *D. serrata* gene sequence to the *D. melanogaster* genome to determine gene orthology using Flybase (Attrill et al. 2016).

## Data Availability Statement

Raw reads for all sequenced lines are available from the NCBI short read archive under Bioproject ID: PRJNA419238. The genomic relatedness matrices used in the population structure analysis are provided in supplementary files (grm_full_Bryc.txt, grm_reduced_Bryc.txt, grm_full_gcta.txt, and grm_reduced_gcta.txt). We have provided the R code, Bryc.R, that implements the test for large eigenvalues (Bryc et al. 2011). The SNP list used to analyse the data in this study is available from Dryad (doi:XXXYYY) and also from www.chenowethlab.org/resources). The CHC phenotype file is provided as the supplementary file pheno.txt. The linear model used to fit the GWAS in ASREML/R model is provided in the file asreml_gwas.R.

## Results and Discussion

### Identification of SNPs

We established a panel of 110 inbred lines of *Drosophila serrata* from wild females caught from a single population in Brisbane Australia and sequenced their genomes. 100 base-pair paired-end reads were mapped to the *Drosophila serrata* reference genome (Allen et al. 2017b) with a mean coverage of 23.5 ± 0.5 reads per line. Using the Joint Genotyper for Inbred Lines (Stone 2012), we identified 15,228,692 biallelic single nucleotide polymorphisms (SNPs) applying a 99% probability threshold. 13,959,239 of these SNPs had a median coverage between 5 and 60 in which over 80% of the lines were genotyped for that variant. Most SNPs segregate at low frequencies (Fig. 1) with 6,090,058 instances of singletons, where a SNP was present in only one line. The majority (62%) of the SNPs were annotated to intergenic regions of the genome while 12% of SNPs annotated to exonic regions and 26% were found to be in introns. A total of 3,709,329 SNPs met the minimum allele frequency threshold (MAF) of 5% to be used for genome-wide association analysis and the estimation of relatedness between lines.

**Figure 1:**
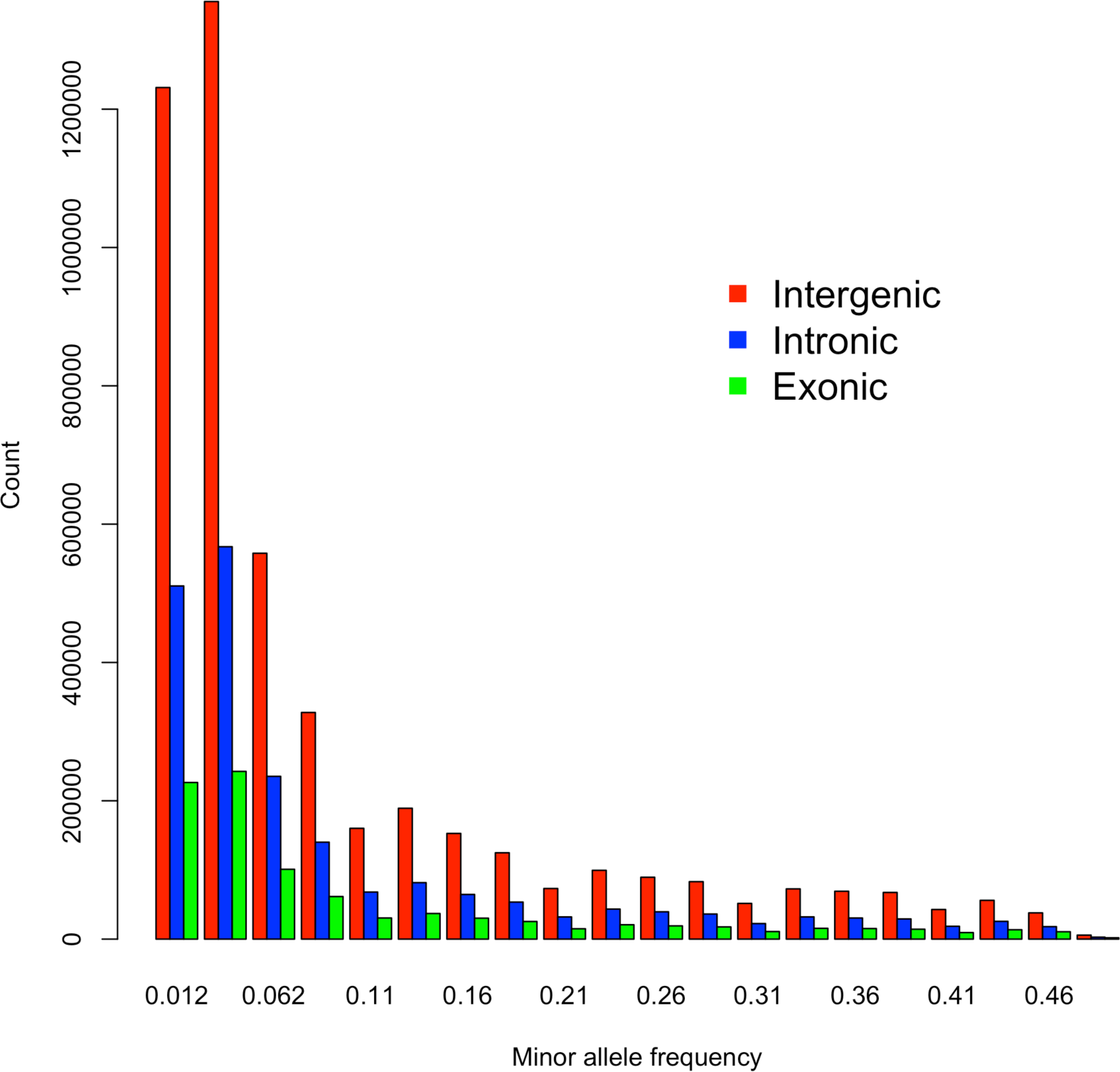
The allele frequency spectrum of SNPs annotated as intergenic (red), intronic (blue), or exonic (green). Singletons (MAF = 0.009) are not shown. We identified 3,748,429 singletons from intergenic regions, 1,535,651 from intronic regions, and 754,075 from exonic regions of the genome.

### Residual heterozygosity

Despite the application of inbreeding for many generations, inbred *Drosophila* lines often contain regions of residual heterozygosity (King et al. 2012; Lack et al. 2015; Mackay et al. 2012; Nuzhdin et al. 1997). After 17-20 generations of inbreeding, residual heterozygosity in our lines was very low and we observed only a small proportion of segregating sites within lines, suggesting that inbreeding had successfully fixed variation across these genomes. Of the 110 inbred lines, 104 had fewer than 2% segregating SNPs and 82 lines had less than 1% segregating SNPs (Fig. 2). Across lines, the average proportion of segregating SNPs was 0.86% ± 0.11%, less than the theoretical expectation of 1.4% for lines that have experienced 20 generations of full-sib mating which corresponds to an expected inbreeding coefficient of F = 0.986 (Falconer and Mackay 1996). This slightly lower than expected level of residual heterozygosity may simply reflect sampling variation, be due to SNPs on the X chromosome, or may indicate purging of partially deleterious alleles during the inbreeding process (Garcia-Dorado 2008; Garcia-Dorado 2012). There was no detectable difference in the fraction of heterozygous SNPs between the lines inbred for 17 and 20 generations (ANOVA: F1,108 = 0.0944, P = 0.76). This result suggests that 17 generations of inbreeding may be sufficient for future line development.

**Figure 2:**
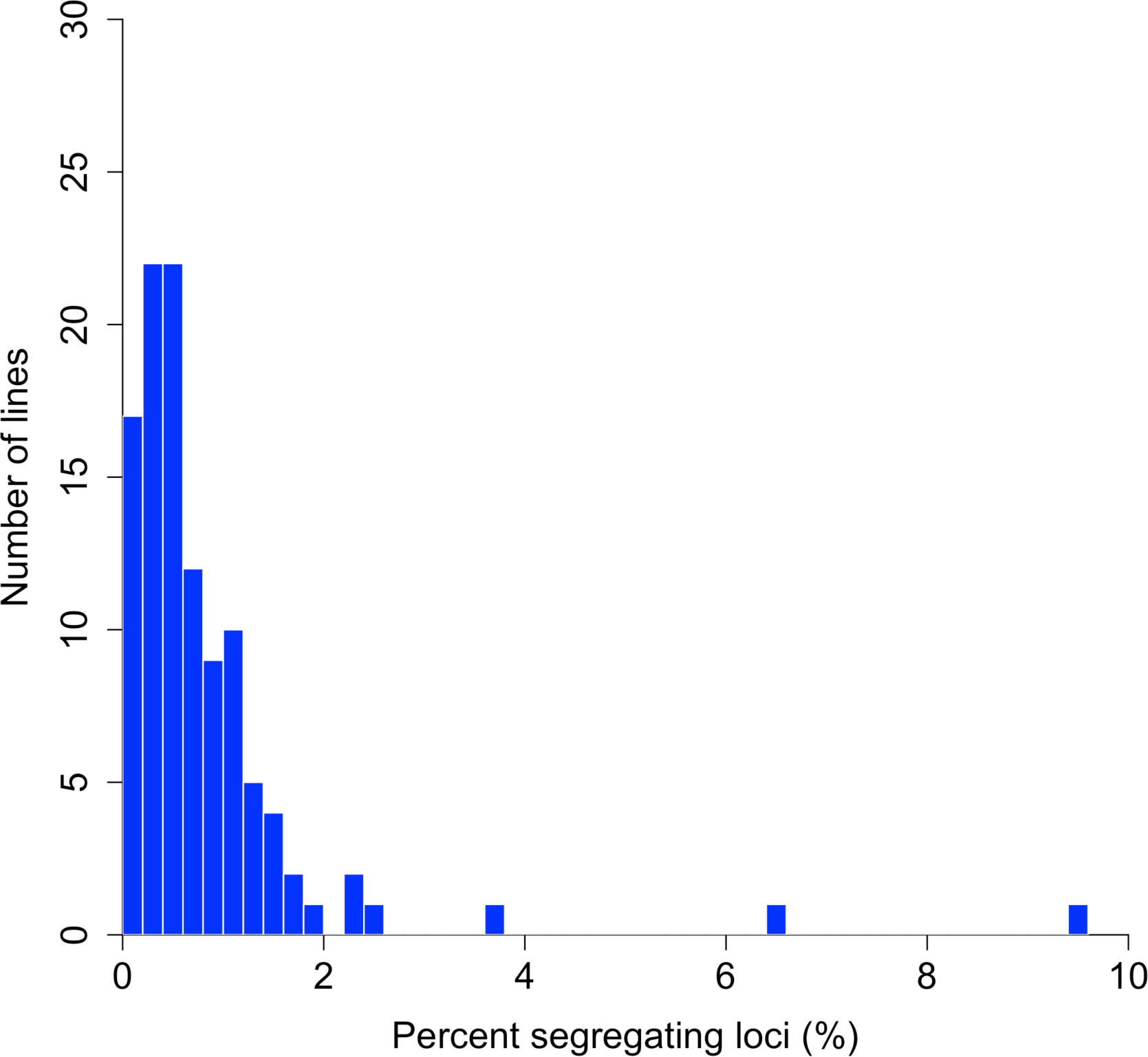
The distribution of residual heterozygosity as measured by the percentage of total genotyped biallelic SNPs (loci) that were called as heterozygous within each inbred line by the Joint Genotyper of Inbred Lines (JGIL).

Although there are several mechanisms that can inhibit the fixation of an allele within an inbred line, when short-read re-sequencing technology is used for genotyping, loci can falsely appear to be segregating due to the presence of paralogous genes or other repetitive DNA sequences (Treangen and Salzberg 2012). If paralogous genes are not represented in the reference genome, DNA sequences from the original gene and a divergent duplicate gene are mapped to the same region of the genome, causing the appearance of segregating loci in the population. When this phenomenon occurs, it is expected that regions of the genome with high “apparent heterozygosity” will associate with a higher read depth than the genome-wide average. Across the genome we found a weak, positive correlation between the level of residual heterozygosity and read depth (Spearman’s ρ = 0.036, p = 2.2 × 10^-16^). Plots of these two factors however, clearly show an alignment of regions with both high levels of residual heterozygosity and read depth, suggesting that the *D. serrata* reference genome could be missing some duplications (Fig. 3). Alternatively, this result could be due to copy number variation among the re-sequenced lines and/or between the reference genome and the 110 lines. We hope that further work and improvement of our genome for this species will elucidate these small regions of residual heterozygosity.

**Figure 3:**
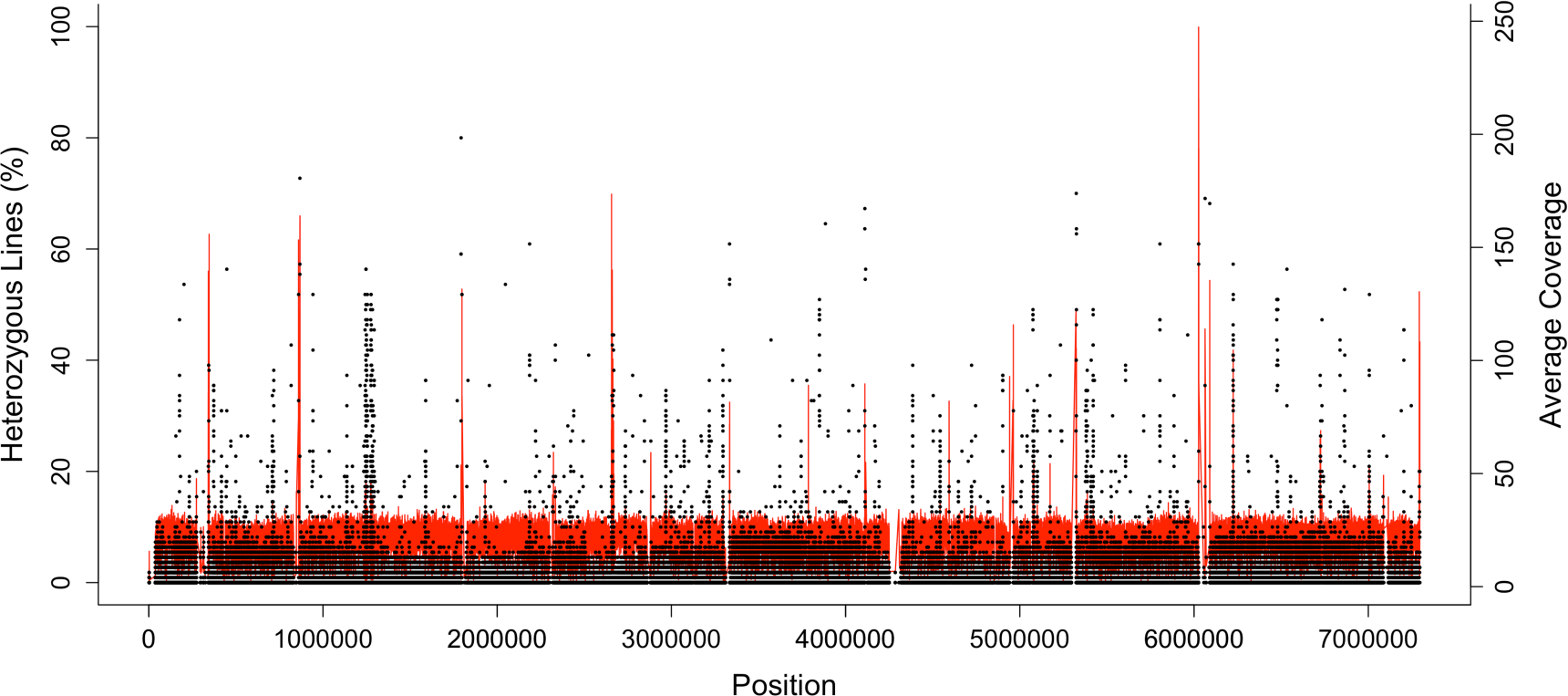
The proportion of lines that are heterozygous at every site along the largest scaffold (scf7180000003208) of the reference genome (black) overlayed with the average read depth at each site shown in red. Peaks in red lines represent potential gene duplications that have collapsed to the same region during assembly.

### Relatedness between lines

Population structure and cryptic relatedness are well known to confound genetic association studies, potentially generating false positive genotype-phenotype associations (Kittles et al. 2002; Knowler et al. 1988). Conceptually, these confounding factors can be described as the unobserved pedigree of the sampled individuals caused by distant relationships (Astle and Balding 2009). The sources of these relationships are varied but include population admixture, inadvertent sampling of close relatives, and the presence of shared chromosomal inversions. Fortunately, the unobserved pedigree can be estimated using marker based approaches. Here we used such an approach to estimate the pairwise relatedness between all lines in the form of a genomic relatedness matrix.

The structure of the genomic relatedness matrix shows that the majority of the DsGRP lines are unrelated as would be expected of a sample from a large, randomly mating population (Fig. 4). We found a pair of lines that were 100% related to one another, most likely due to contamination either during the inbreeding process or DNA extraction and subsequent library preparation. Generally however, this panel of flies exhibits a lower level of relatedness compared to the DGRP, where 2.7% of the 20,910 possible pairs of lines had estimates of pairwise relatedness over 0.05 (Huang et al. 2014) compared to 0.08% of 5,995 pairs of lines reported here. The discrepancy between the levels of relatedness between the DsGRP and DGRP is potentially due to the different demographic histories of the founding populations that generate population structure. North American populations of *D. melanogaster* have relatively complex demographic histories with admixture of African and European ancestors and instances of secondary contact (Pool et al. 2012) compared to the endemic population of *D. serrata*.

**Figure 4:**
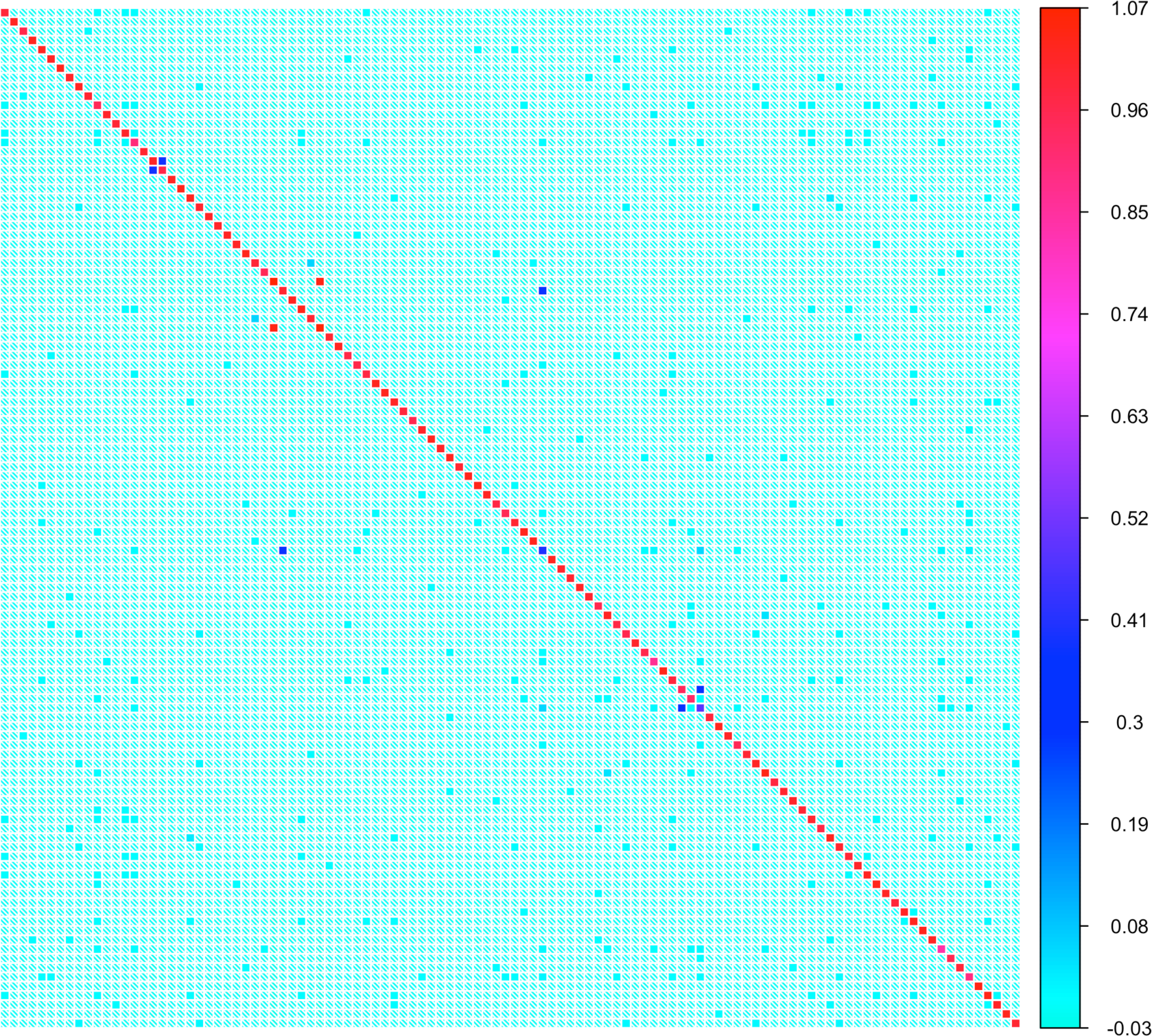
Heat-map of the genomic relatedness matrix for each pair of the 110 inbred DsGRP lines. Each coloured squared is an estimate of genomic relatedness between a pair of inbred lines estimated by GCTA (Yang et al. 2011). The shade of colour represents the degree of relatedness, with light blue showing low levels of relatedness and red high levels of relatedness.

Another likely explanation for the increased levels of relatedness in the DGRP is the presence of common segregating inversions. While chromosomal inversions are known to segregate in *D. serrata*, their frequency and number tends to increase in populations approaching the equator (Stocker et al. 2004). Therefore, it may be the case that founding the DsGRP from the higher latitude of Brisbane has resulted in sampling relatively few inversions. As of yet, these lines have not been karyotyped; however the low levels of relatedness and the lack of any bimodal distribution for residual heterozyosity, such as the one found in the DGRP (Huang et al. 2014), where a portion of the lines had high levels of segregating SNP loci (15-20%), suggests that segregating inversions are negligible in this population.

We performed an eigendecomposition of the genomic relatedness matrix to test for the presence of population structure using the approach outlined in Bryc et al. (2013). This analysis revealed two large eigenvalues (*Λ*_1_ = 20.08 and *Λ*_2_ = 1.12) that were greater than that expected for a random relatedness matrix of equal size (Threshold = 0.993) (Fig. 5). There is therefore evidence that the full set of 110 of lines contain substructure in the form of two subpopulations. We reasoned that the second large eigenvalue was likely caused by the four pairs of lines that were highly related to each other (*A_jk_* = 0.29, 0.38, 0.39, and 1.04; Fig. 4). To test this, we repeated the analysis after randomly removing one line from each of the four pairs of closely related lines. Confirming the prediction, there was only one significantly large eigenvalue in this second analysis (*Λ*_1_ = 19.66). Such a result is expected when the data includes only a single population. To summarise, after the four highly-related lines have been removed, there is no clear evidence for population structure in the DsGRP that would lead to spurious genotype-phenotype associations in genome-wide association analysis.

**Figure 5:**
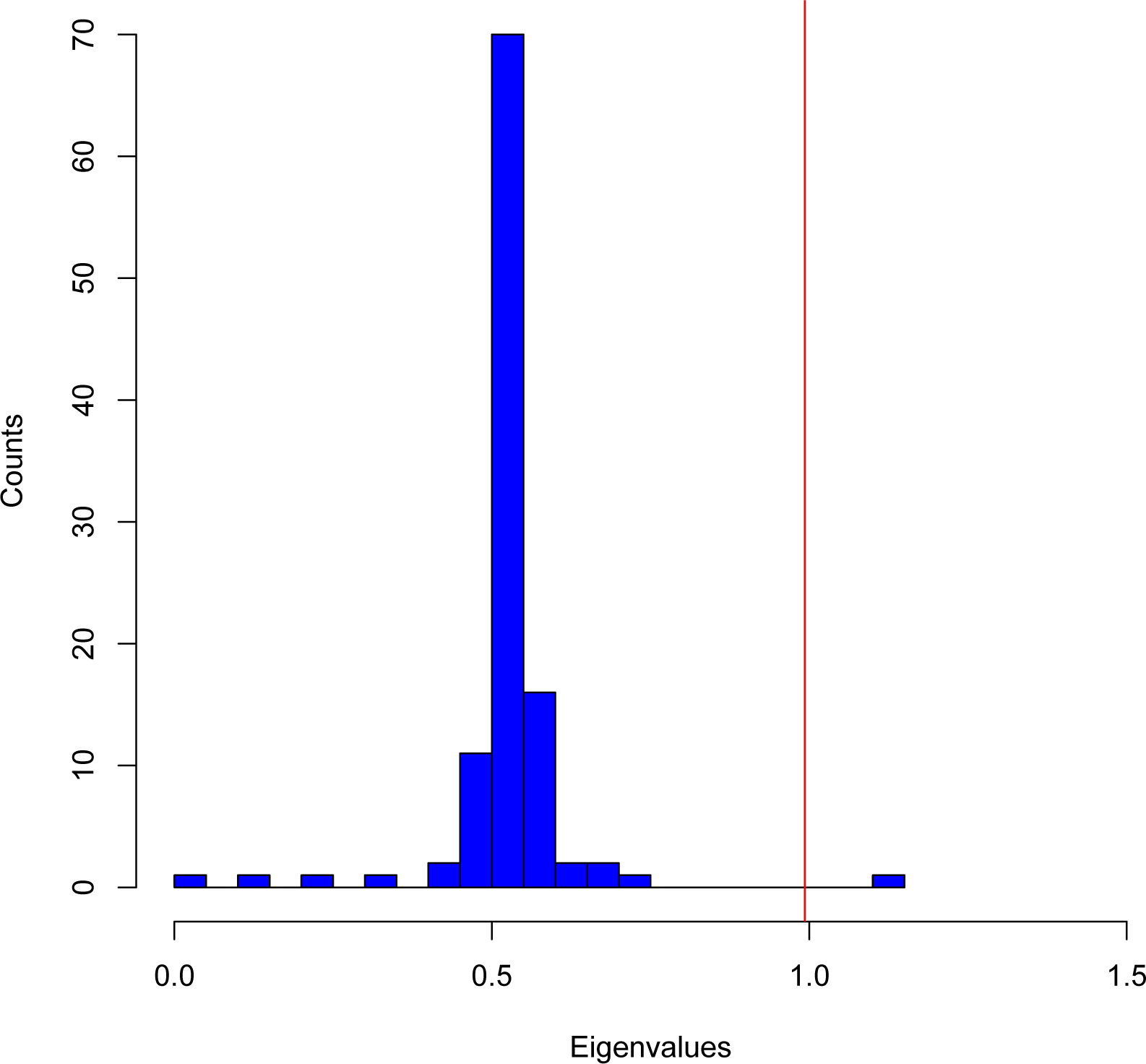
Distribution of eigenvalues from an eigendecomposition of the genomic relatedness matrix for all 110 lines excluding one large eigenvalue where *Λ* = 20.08. Eigendecomposition of the genomic relatedness matrix, **X**, which was scaled by equation 2.8 in Bryc et al. (2013) using values of N= 3,709,328 and M=110. Here, N corresponds to the number of SNPs used to estimate **X** and M is the number of lines. Only SNPs with without missing data were used for this analysis. The red vertical line corresponds to the significance threshold, *t*, for declaring an eigenvalue larger than that expected for a random relatedness matrix. *t* = (1 + *F*)/2 and corresponds to a value of 0.993 with our expected inbreeding coefficient (*F*) of 0.986 after 20 generations of full-sib mating. The two largest eigenvalues were significant, *Λ*_1_ = 20.08 (not plotted) and *Λ*_2_ = 1.12.

### Linkage Disequilibrium

The rapid decay of linkage disequilibrium with genomic distance is a common feature of *Drosophila* species with *r*^2^ dropping below 0.1 within 100 base pairs (Long et al. 1998; Mackay et al. 2012). This allows for higher resolution mapping compared to other species such as maize and humans in which the equivalent decay does not occur until after approximately 2000 base pairs (Remington et al. 2001) and 50,000 base pairs (Koch et al. 2013), respectively. In the DsGRP, linkage disequilibrium decays rapidly with r^2^, on average, dropping below 0.1 after 75 base pairs. Surprisingly, we observe faster decay on the X chromosome compared to the autosomes (Fig. 6), contrary to Mackay et al. (2012), despite the fact that the X chromosome has a smaller effective population size than the autosomes.

**Figure 6:**
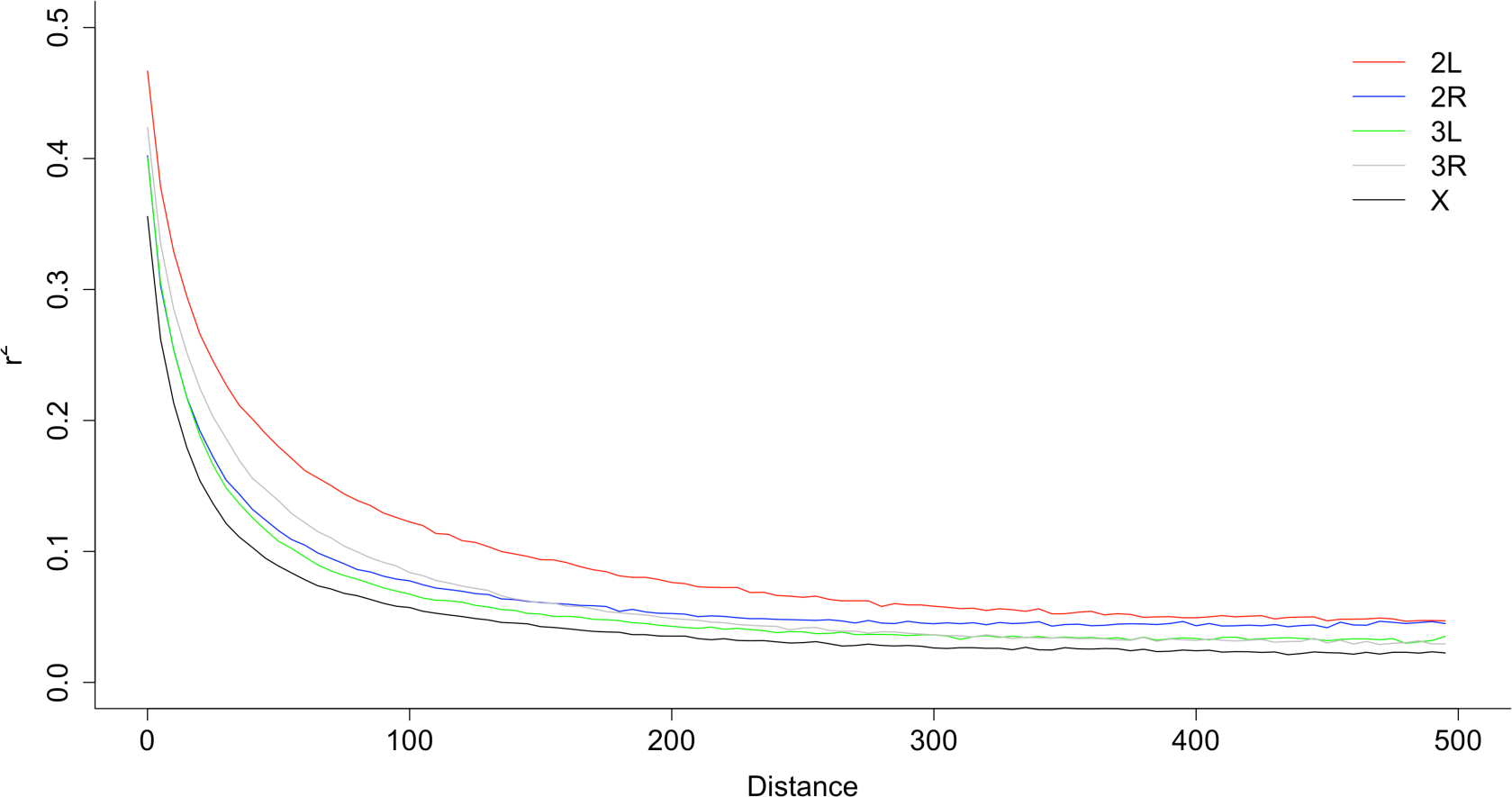
Decay of linkage disequilibrium (r^2^) between SNPs with genomic distance (bp) in the DsGRP. Values are averaged across each chromosome.

### Nucleotide diversity and neutrality

For the two cosmopolitan species of *Drosophila* that have been studied extensively, *D. melanogaster* and *D. simulans*, the ancestral populations from Africa consistently exhibit higher levels of polymorphism compared to the derived populations from America and Europe (Andolfatto 2001; Baudry et al. 2004; Begun and Aquadro 1993; Grenier et al. 2015; Lack et al. 2015). Presumably, nucleotide diversity is reduced during bottleneck events associated with the colonisation of new habitat. The relatively high estimates of nucleotide diversity for *D. mauritiana*, an endemic species from Mauritius, bolster this trend (Garrigan et al. 2014). We therefore expected that our population of *D. serrata*, founded from the species’ ancestral range, would exhibit relatively high levels of nucleotide diversity. We estimated nucleotide diversity (π) along the major chromosome arms 2L, 2R, 3L, 3R, and X using a 50 kilobase non-overlapping sliding window approach (Table 1, Fig. 7). Averaged across the genome, we estimated that π = 0.0079, which is consistent with the pattern seen in other species of relatively increased levels of nucleotide diversity for populations from ancestral ranges compared to more recently established population outside of the ancestral range (Andolfatto 2001; Baudry et al. 2004; Begun and Aquadro 1993; Grenier et al. 2015; Lack et al. 2015).

**Figure 7:**
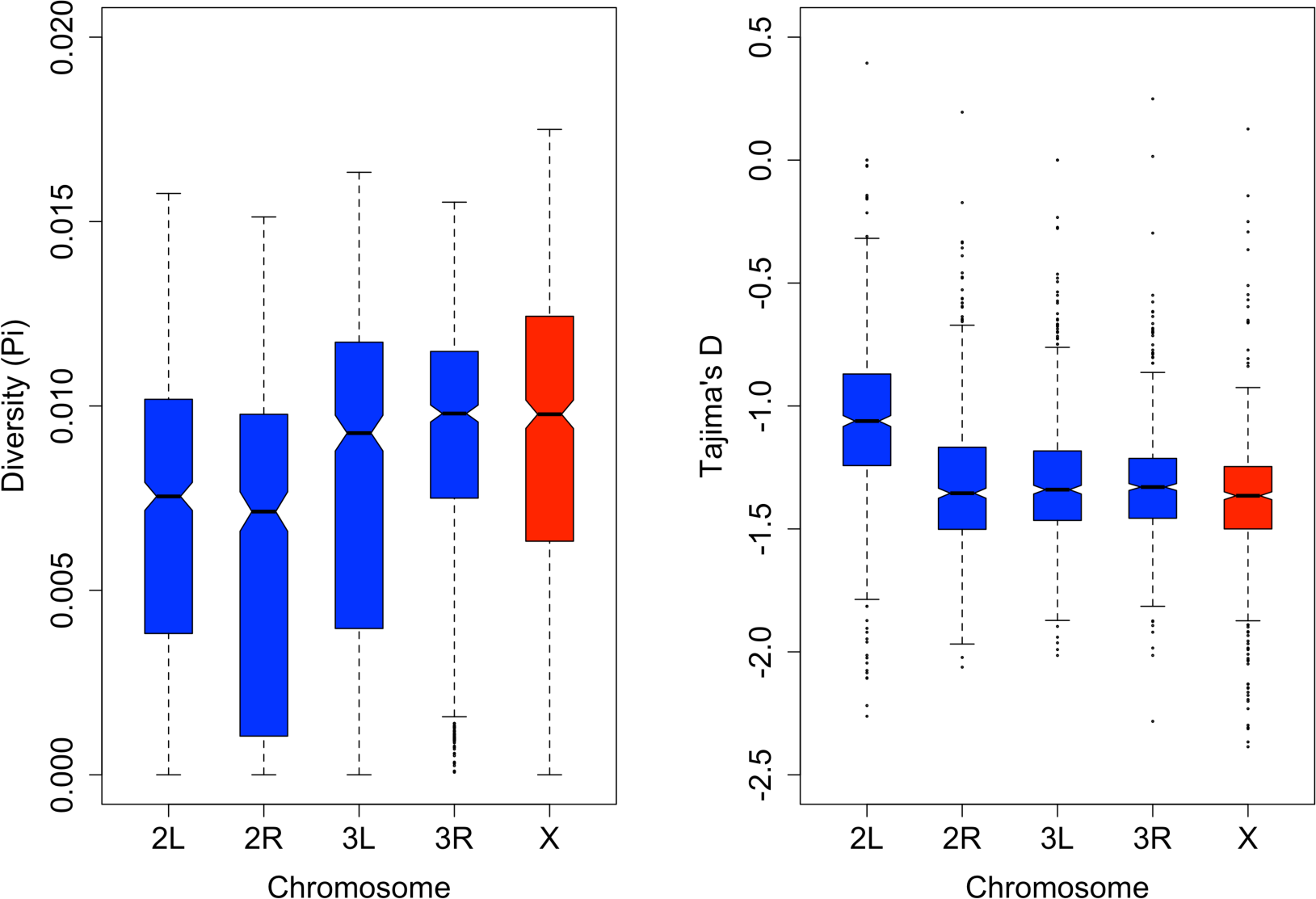
Boxplots of nucleotide diversity and Tajima’s D by chromosome arm. Shown are the estimates from 50 kilobase non-overlapping sliding windows with the breakdown of the number of windows per chromosome as follows: 2L = 691, 2R = 664, 3L = 641, 3R = 741, and X = 628.

Using the same sliding window approach, we tested for departures from neutrality using Tajima’s D (Table 1, Fig. 7) (Tajima 1989). As with other populations of *Drosophila*, the DGRP (Mackay et al. 2012), the Zimbabwean population in the Global Diversity Lines (Grenier et al. 2015), and *D. mauritiana* (Garrigan et al. 2014), Tajima’s D was negative across the entire genome (Tajima’s D = -1.27). This is consistent with an abundance of rare alleles and could be indicative of population expansion or the occurrence of selective sweeps, however this statistic cannot distinguish the effects of demography from selection. In the DsGRP, chromosome arm 2L has higher estimates of Tajima’s D compared to the other chromosome arms (Fig. 7). The causes for this pattern will hopefully be resolved through a more in-depth population genomic analysis, as chromosome 2L may potentially harbour more genomic regions that experience balancing selection, which would increase the average value of Tajima’s D.

### Genome-wide association analysis of female CHC expression

A major motivation for developing the DsGRP was to begin connecting molecular variation with standing variation for some of the well-studied quantitative traits of *D. serrata*. Here, we applied genome-wide association analysis to the expression of the CHC 2-Me-C _28_in females and identify new candidate genes that might influence trait variation. Using our mixed model approach, we found 4 SNPs that passed the 0.05 significance threshold after Bonferroni multiple test correction (Figure 8). Two of the SNPs are situated within genes, while the other two lie within 3kb of genes. We found a further 189 SNPs with p-values lower than a suggestive threshold of 10^-5^.

**Figure 8:**
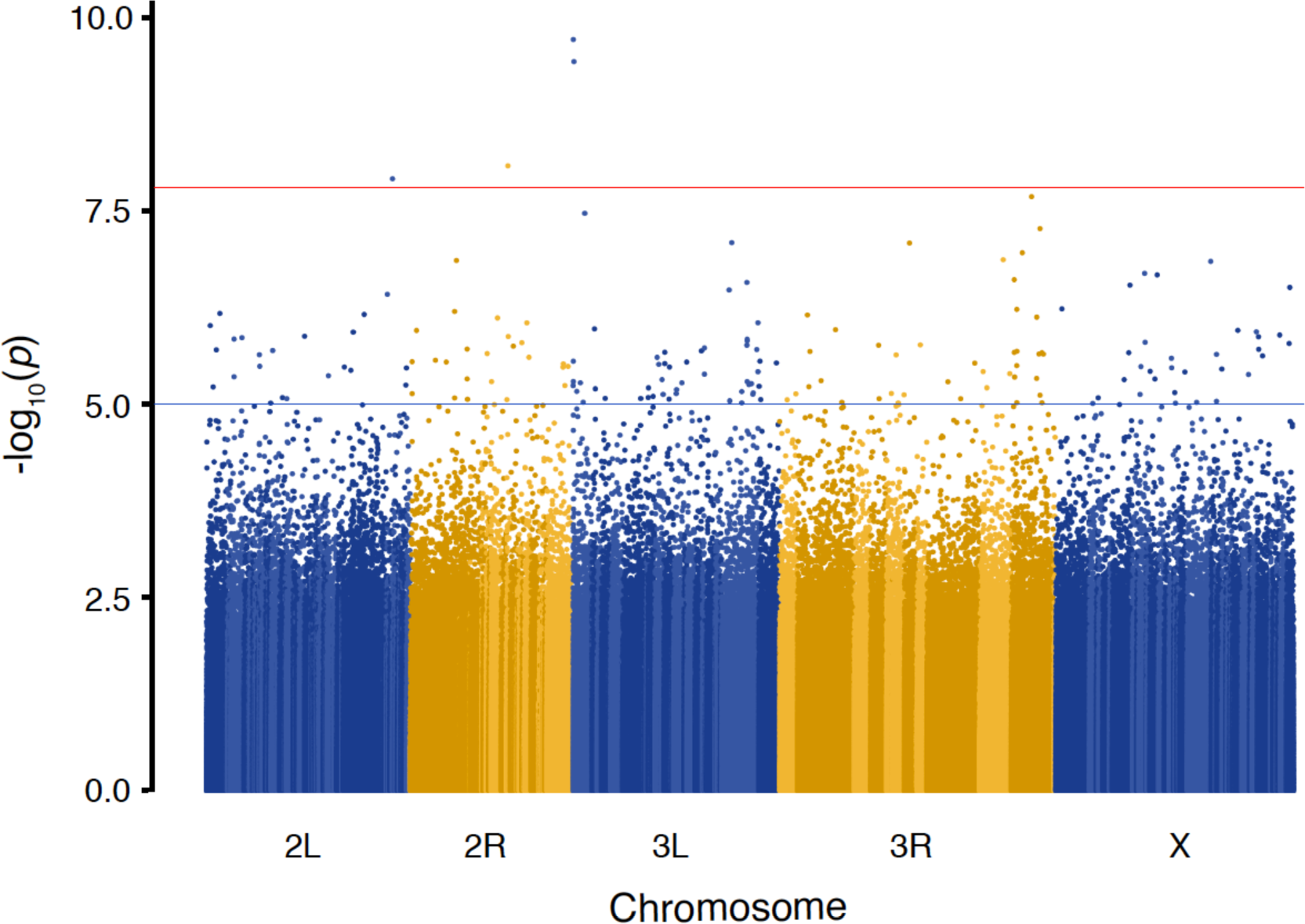
Genome-wide association for 2-Me-C _28_expression in female *Drosophila serrata.* Red line indicates Bonferroni threshold corresponding to P = 0.05 and the blue indicates an arbitrary significance threshold of P = 10^-5^. A total of 3,318,503 biallelic SNPs were analyzed with a minimum minor allele frequency of 0.5. Alternating colour shading within chromosomes indicates different contigs in the *D. serrata* genome assembly. Contig order has not yet been established for the genome.

We note that a GWAS performed on line mean data using the GCTA program (Yang et al. 2011), which like many other mixed model GWAS applications, estimates the polygenic variance only once, detected no SNPs above Bonferroni threshold and only 34 under the arbitrary threshold of 10^-5^. We also applied our ASREML approach to line means rather than individuals and found that it detected exactly the same 4 SNPs above Bonferroni as our approach in eq. 2 (190 SNPs had p-values lower than 10^-5^). It therefore appears that the increase in detection rate is in this case mainly due to the ASREML model re-estimating the polygenic variance for each SNP tested which results in an exact, rather than approximate, calculation of the test statistic (Zhou and Stephens 2012).

The majority of the literature regarding the expression of CHCs has identified genes that are related to their production within specialised cells, oenocytes. These genes constitute the major biosynthetic pathway known for CHC production and are involved with fatty-acid synthesis, elongation, desaturation, and reduction (Chertemps et al. 2007; Chertemps et al. 2006; Chung et al. 2014; Fang et al. 2009; Labeur et al. 2002; Marcillac et al. 2005; Wicker-Thomas et al. 2015). Although none of the genes associated with the statistically significant SNPs found in this study are involved in this biosynthetic pathway, there are other biological processes involved with CHC expression, as measured by hexane washes from the cuticle. How CHCs are transported from the oenocytes to the cuticle is unknown, this study provides a potential candidate gene involved in this process. One of the significant SNPs resides in the gene *Cht9*, a *chitinase* found on chromosome 2R. *Cht9*, along with a number of other *chitinases* and *imaginal-disc-growth-factors* are important for the development of epithelial apical extracellular matrix, which controls the development and maintenance of wound healing, cell signalling, and organ morphogenesis in *Drosophila* (Galko and Krasnow 2004; Turner 2009). Knocking out the expression of *Cht9* with RNAi leads to deformed cuticles, inability to heal wounds, and defects in larval and adult molting (Pesch et al. 2016), and here, we provide evidence that variation in this gene may also influence other cuticular traits such as CHC abundance. Notwithstanding the small number of lines, the genome-wide association analysis presented here, combined with a previous study that identified the major role of the transcription factor *POU domain motif 3* (*pdm3*) for polymorphic female-limited abdominal pigmentation (Yassin et al. 2016), illustrate the potential of the DsGRP to discover novel regions of the genome that underpin the genetic architecture of traits.

## Conclusion

We have assembled a new resource for the study of quantitative traits and population genomic variation in a non-model *Drosophila* species within its endemic distribution. These reproducible genotypes sampled from a single population not only provide a rich genomic dataset suitable for population genomic studies, but also provide a critical resource for the discovery of genetic variants underlying ecologically important quantitative traits. We hope that the DsGRP will provide a useful complement to other *Drosophila* resources such as the DGRP (Mackay et al. 2012), the DSPR (King et al. 2012), and the Drosophila Genome Nexus (Lack et al. 2015).

In this first characterisation of the DsGRP at the genomic level, we have shown that the inbreeding process has been successful in homogenising the majority of the genome of each of the lines. Through the estimation of the genomic relatedness matrix we have shown that the DsGRP represents a random sample from a large population that contains very low levels of cryptic relatedness. These characteristics, along with rapid decay of linkage disequilibrium, make the DsGRP an ideal resource for the application of genome-wide association analysis and for generating new multifounder QTL mapping populations that will boost mapping power.

## Acknowledgements

We thank EK Delaney for comments on the manuscript and NC Appleton for assistance in the laboratory. This work was supported by an Australian Postgraduate Award to ARR and funds from the Australian Research Council and The University of Queensland awarded to SFC.

